# Enzyme specific isotope effects of the Nap and Nar nitrate reductases

**DOI:** 10.1101/2020.11.17.386888

**Authors:** Ciara K. Asamoto, Kaitlin R. Rempfert, Victoria H. Luu, Adam D. Younkin, Sebastian H. Kopf

## Abstract

Dissimilatory nitrate reduction (DNR) to nitrite is the first step in denitrification, the main process through which bioavailable nitrogen is removed from ecosystems. DNR fractionates the stable isotopes of nitrogen (^14^N, ^15^N) and oxygen (^16^O, ^18^O) and thus imparts an isotopic signature on residual pools of nitrate in many environments. Data on the relationship between the resulting isotopic pattern in oxygen versus nitrogen isotopes (^18^ε / ^15^ε) suggests systematic differences exist between marine and terrestrial ecosystems that are not fully understood. DNR can be catalyzed by both cytosolic (Nar) and periplasmic (Nap) nitrate reductases, and previous work has revealed differences in their ^18^ε / ^15^ε isotopic signatures. In this study, we thus examine the ^18^ε / ^15^ε of six different nitrate-reducing microorganisms that encode Nar, Nap or both enzymes, as well gene deletion mutants of the enzymes’ catalytic subunits (NarG and NapA) to test the hypothesis that enzymatic differences alone could explain the environmental observations. We find that the distribution of the ^18^ε / ^15^ε fractionation ratios of all examined nitrate reductases form two distinct, non-overlapping peaks centered around a ^18^ε / ^15^ε proportionality of 0.55 and a ^18^ε / ^15^ε proportionality of 0.91, respectively. All Nap reductases studied to date cluster around the lower proportionality (0.55) and none exceed a ^18^ε / ^15^ε proportionality of 0.68. Almost all Nar reductases, on the contrary, cluster tightly around the higher proportionality (0.91) with no values below a ^18^ε / ^15^ε proportionality of 0.84 with the notable exception of the Nar reductases from the genus *Bacillus* which fall around 0.62 and thus closely resemble the isotopic fingerprints of the Nap reductases. Our findings confirm the existence of two remarkably distinct isotopic end-members in the dissimilatory nitrate reductases that could indeed explain differences in coupled N and O isotope fractionation between marine and terrestrial systems, and almost but not fully match reductase phylogeny.

## Introduction

Nitrogen is an essential nutrient for life and consequently the availability of nitrogen is a vital control on ecosystem productivity. Anthropogenic activity has severely altered the natural balance of the nitrogen cycle. In particular, the use of the Haber-Bosch reaction to synthesize fertilizers has resulted in excess amounts of nitrate and ammonium being introduced into ecosystems ^1,2^. Assessing the outcomes of excess nitrogen inputs into ecosystems requires a mechanistic understanding of the competing processes that affect nitrogen cycling in the environment.

Fig. 1A highlights key reductive and oxidative steps in the nitrogen cycle, all of which are catalyzed by microorganisms ^3^. The enzymes bacteria use to reduce or oxidize nitrogen intermediates in the nitrogen cycle impart a kinetic isotope effect on the stable isotopes of nitrogen (^14^N, ^15^N) and oxygen (^16^O, ^18^O) ^4–9^. Because nitrogen fixation by most nitrogenases does not impart strong isotopic fractionation ^9–13^, redox cycling of fixed nitrogen, especially the isotopic fractionation associated with dissimilatory nitrate reduction to nitrite, controls the isotopic composition of bioavailable nitrate in many environmental systems. Dissimilatory nitrate reduction is the first step for two processes in the nitrogen cycle, denitrification to N_2_ and dissimilatory nitrate reduction to ammonium (DNRA, also referred to as nitrate ammonification) (Fig. 1A). Although these processes serve different roles, both impact the isotopic composition of residual nitrate in ecosystems through the nitrate reduction step.

**Fig. 1.**
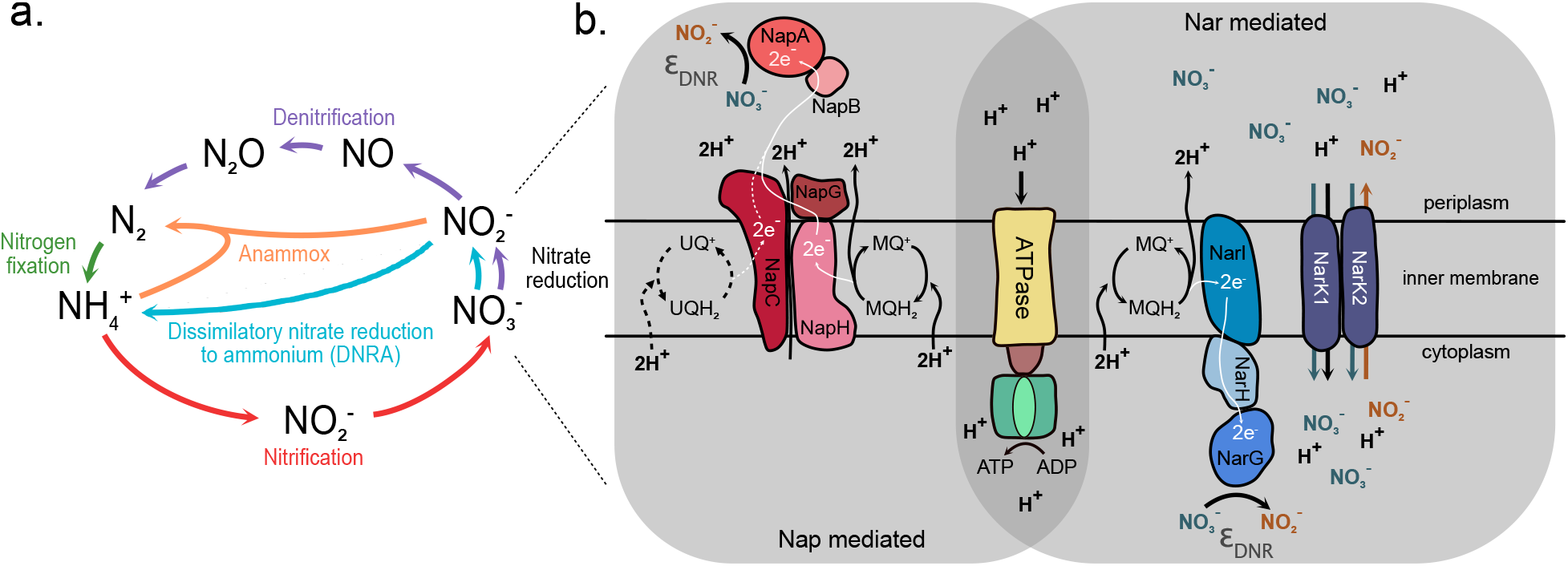
(Top left) An overview of the nitrogen cycle with focus on the dissimilatory nitrate reduction step. (Right) The schematic highlights differences for how nitrate reduction is catalyzed in Nap versus Nar enzymes. Isotope fractionation (ε_DNR_) occurs during the reduction of nitrate to nitrite. White lines indicate the direction of electron transfer. Black lines indicate proton translocation. In the case of Nap reductases, there are two main potential pathways for nitrate reduction to occur. Bacteria may express NapABC (dashed lines), where NapC oxidizes ubiquinol (UQH_2_) to ubiquinone (UQ^+^), liberating two protons and two electrons. The electrons are transferred to NapB then NapA. Alternatively, a bacterium may express NapABCGH (solid lines). Here NapH oxidizes menaquinol (MQH_2_) to menaquinone (MQ^+^) and the electrons have an additional transfer step from NapG to NapC, translocating two additional protons. The Nar reductase uses NarI to oxidize UQH_2_ to UQ^+^ and transfers electrons to NarH then NarG. NarK1 is a symporter that transports nitrate into the cytoplasm with a proton. NarK2 is an antiporter that couples the import of nitrate to the export of nitrite.

The proportionality of N and O isotope fractionation (^18^ε / ^15^ε) associated with nitrate reduction in marine ecosystems generally follows a proportionality of 0.9 to 1.0 ^14–19^. In terrestrial ecosystems, observational data with coupled N and O isotope measurements is more limited (summarized in Fig. 2) but the existing data suggests that the ^18^ε / ^15^ε proportionality covers a broader and generally lower range of values between 0.5 to 0.7 ^20–26^. To date, these systematic differences in ^18^ε / ^15^ε proportionality are not fully understood and may indicate that we are missing a key feature about how nitrogen cycling processes create the isotopic signatures of nitrate observed in nature. Biogeochemical modelling and recent culturing work suggest that the terrestrial observations of low ^18^ε / ^15^ε values could be the result of oxidative overprinting of the isotopic signal of nitrate reduction by a combination of nitrate producing processes such as anaerobic ammonium oxidation (annamox), nitrification, and enzymatic reversibility during nitrate reduction ^8,27^. However, an alternative hypothesis first proposed by Granger *et al.* (2008) suggests that differences in the ^18^ε / ^15^ε proportionality observed in nature could actually be a consequence of enzymatic differences during nitrate reduction.

**Fig. 2.**
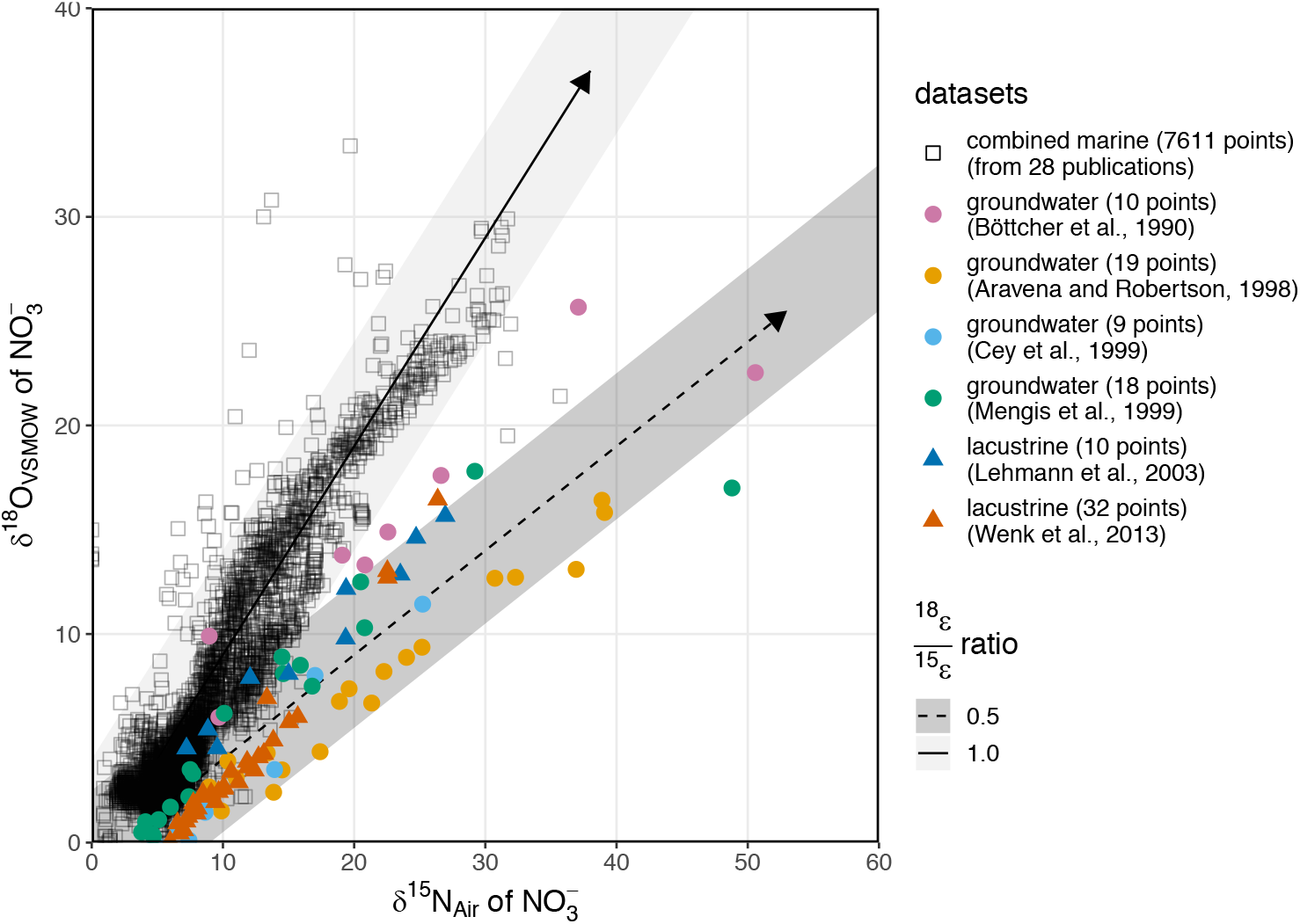
A compilation of nitrate isotopic data collected from environmental samples subset into marine and terrestrial/ freshwater ecosystems. Solid lines and dashed lines indicate ^18^ε / ^15^ε proportionalities of 1.0 and 0.5, respectively, with gray shaded bands showing a range of possible intercepts. See SI for details on the literature data.

Dissimilatory nitrate reduction can be catalyzed by the periplasmic enzyme Nap (catalytic subunit NapA) and the membrane bound cytosolic enzyme Nar (catalytic subunit narG). Bacteria can harbor either or both of these nitrate reductases ^28–30^ and neither is linked exclusively to either denitrification or DNRA. The few studies that have specifically examined the isotope effects of Nap reductases ^7,31,32^ indicate that Nap N isotope fractionation (^15^ε) ranges from 11.4-39.8‰, overlapping with that of Nar reductases (6.6-31.6‰). However, the proportionality between O and N isotope fractionation appears to differ between Nap and Nar-based nitrate reduction. The purple photoheterotroph *Rhodobacter sphaeroides* and the chemotrophic sulfur oxidizer *Sulfurimonas gotlandica* both have only a Nap reductase and were examined by Granger et al. (2008); Treibergs & Granger (2017) and Frey et al., 2014, respectively. The isotopic data from the Nap reductases in these organisms revealed ^18^ε / ^15^ε values between 0.57 – 0.68 for *R. sphaeroides* and 0.43 – 0.68 for *S. gotlandica,* in contrast with the ^18^ε / ^15^ε proportionality of ~0.9 in Nar based nitrate reduction ^7,32–34^. Here, we present experimental results based on six different nitrate-reducing microorganisms that encode Nar, Nap or both enzymes, as well gene deletion mutants of the enzymes’ catalytic subunits (NarG and NapA) to test the hypothesis that differences in ^18^ε / ^15^ε proportionality may stem solely from enzymatic differences and explore the implications of our results for the environmental interpretation of nitrate isotope signatures.

## Methods

### Strains

All strains cultured for this study have either the gene for the cytosolic nitrate reductase (narG), the gene for the periplasmic nitrate reductase (napA), or both. The strains that have both narG and napA are *Pseudomonas aeruginosa* PA14 (DSM 19882) and *Paracoccus denitrificans* PD1222, a derivative of DSM 413 ^35,36^. The strains with only napA are *Desulfovibrio desulfuricans* DSM 642, *Shewanella loihica* DSM (17748) ^37–39^, and a markerless narG deletion mutant of *P. aeruginosa* PA14 ^40^, hereafter referred to as PA14 Δnar. The strains with only narG are *Bacillus vireti* (DSM 15602), *Bacillus bataviensis* (DSM 15601) ^41^, and a markerless napA deletion mutant of *P. aeruginosa* PA14, hereafter referred to as PA14 Δnap.

### Culturing

PA14 strains were grown at 30°C and 37°C (PA14 Δnar) in defined MOPS minimal media amended with 25mM sodium succinate as the sole carbon source ^42^, as well as 25g/L LB broth. *B. vireti* and *B. bataviensis* were grown at 30°C in 30g/L tryptic soy broth (TSB) amended with 13mM glucose and 11mM sodium succinate ^43^. *D. desulfuricans* was grown at 30°C in Postgate’s defined medium ^44^, which contains 20mM lactate and 1 g/L yeast extract as carbon sources as well as sodium thioglycolate (0.1g/ L) as a reductant. *S. loihica* was grown at 30°C in a phosphate buffered minimal salts medium amended with 5, 25, or 30mM sodium lactate as the sole carbon source (Yoon et al. 2015). *P. denitrificans* was grown at 30°C in a defined minimal salts medium amended with 25 mM sodium acetate as the sole carbon source (Hahnke et al. 2014). For all nitrate reduction experiments, NaNO_3_^−^was injected from a concentrated stock solution into each culture tube. *S. loihica* media was amended with approximately10mM NaNO_3_ in this way and all other media recipes were amended with approximately 25mM NaNO_3_. Exact concentrations in each sample were confirmed by ion chromatography.

For all anaerobic growth experiments, media was sparged with N_2_ gas and cultures were incubated while shaking at 250 rpm in balch tubes containing 20mL of media and 5mL of N_2_ headspace at 1.1bar and sealed with blue butyl rubber stoppers. For aerobic growth, culture tubes were incubated while shaking at 250rpm. Agar plates for reviving strains from frozen stock were prepared by amending each media recipe with 15 g/L agar. All strains except *D. desulfuricans* (an obligate anaerobe) were revived on aerobic agar plates and passaged three times in liquid medium before inoculating isotope fractionation experiments with 1% culture (v/v). *D. desulfuricans* was inoculated directly from freezer stocks into anaerobic culture medium and passaged 5 times before inoculating isotope fractionation experiments.

### Isotope Fractionation Experiments

All strains were grown in triplicate in their respective media in the presence of nitrate and sampled at regular intervals for nitrate consumption and nitrate isotopic composition. Growth was monitored directly in the culture tubes by optical density (OD) using a Spectronic 20 spectrophotometer at a wavelength of 660nm for *B. vireti* and *B. bataviensis*, and 600nm for all other strains. At each time point, approximately 2mL of sample was withdrawn through the stopper using a 23-gauge needle attached to a syringe. Syringes were flushed with nitrogen prior to sampling to preserve the anaerobic environment within the balch tubes. Samples were filter sterilized with 0.2μm PES filters, aliquoted for later quantification and isotopic analysis, and stored at −20°C. Aliquots for ion chromatography (IC) were immediately diluted in 0.1M NaOH (pH 11) to stabilize nitrite. For *P. aeruginosa* and *B. vireti*, the experiment was additionally repeated in media made from ^18^O enriched water (OLM-240-10-1, Cambridge Isotope Laboratories, Inc.) at a final δ^18^O_water_ of approximately +100‰.

### Sample Analysis

Nitrate and nitrite concentrations were quantified using a Dionex ICS-6000 Ion Chromatograph equipped with an IonPac AS11-HC column and a variable wavelength absorbance (UV/Vis) detector to allow for accurate analyte detection in complex media (LB, TSB). Samples were eluted isocratically with 20mM KOH at a flow rate of 1mL/ minute. Nitrate and nitrite peaks were measured at a wavelength of 210nm and quantified against laboratory standards prepared in the same media backgrounds. The N and O isotopic composition of nitrate was determined in the Sigman Lab at Princeton University using the denitrifier method ^47,48^ with 20 nmol nitrate per analysis. Nitrite removal was performed prior to isotopic analysis for all samples with nitrite concentrations > 1% nitrate using the sulfamic acid method ^49^. The isotopic measurements were calibrated against the potassium nitrate reference standards IAEA-NO3 (δ^15^N = 4.7‰ vs. air, δ^18^O = 25.6‰ vs. Vienna Standard Mean Ocean Water (VSMOW)), provided by the International Atomic Energy Agency and USGS34 (δ^15^N = −1.8‰ vs. air, δ^18^O = −27.9‰ vs. VSMOW) provided by the United States Geological Survey, each measured at two different concentrations every 8 samples to correct for injection volumes. Analytical runs were corrected for instrument drift based on an N_2_O drift monitoring standard. All isotopic data are reported in conventional delta notation versus the international reference scales for N (Air) and O (VSMOW): δ^15^N = ([^15^N/^14^N]_sample_/[^15^N/^14^N]_air_ – 1) and δ^18^O = ([^18^O/^16^O]_sample_/[^18^O/^16^O]_VSMOW_ – 1). δ values reported in per mil (‰) are implicitly multiplied by a factor of 1000 ^50^. The analytical precision of the nitrate monitoring standard used across all analytical runs was 0.06‰ for δ^15^N and 0.69‰ for δ^18^O (1 *σ*, n=33). Additionally, all fractionation experiments were run using the same nitrate source in different media, thus initial time points across all experiments provide an estimate of sample analytical precision: 0.07‰ for δ^15^N and 0.43‰ for δ^18^O (1 σ, n=52).

### Calculations

#### Isotope effects

the nitrate δ^15^N and δ^18^O measurements were fit to the following linear equations to estimate the N and O isotope effects (^15^ε and ^18^ε) and isotope effect proportionality (^18^ε / ^15^ε) imparted on nitrate during microbial nitrate reduction from the slope of the regressions ^51^:

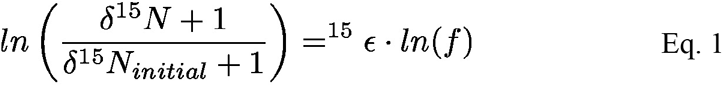

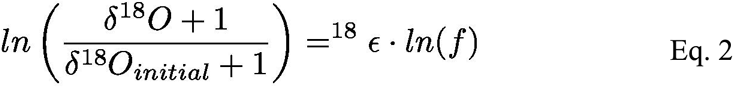

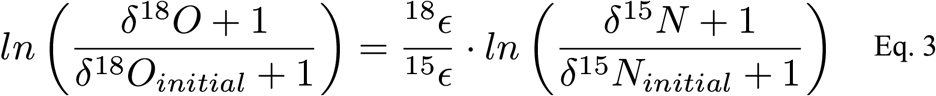

where *f* = [NO_3_^−^]/[NO ^−^] is the fraction of nitrate remaining and δ and ε values in per mil (‰) are implicitly multiplied by a factor of 1000 ^50^. The errors of the regression slopes were used to estimate standard errors for ^15^ε (Eq. 1), ^18^ε (Eq. 2), and ^18^ε / ^15^ε (Eq. 3). Note that for this implementation of the Rayleigh distillation model (Eq. 1 & 2), normal kinetic isotope effects (reflecting higher reaction rates of the lighter isotopes) are negative (ε < 0) and are reported as such in Table S1. The opposite convention with normal kinetic isotope effects reported as ε > 0 is also not uncommon and all comparisons with literature data carefully consider the convention used in each publication. For visual representation of Eq. 3 in figures, the following more intuitive but slightly less accurate linearizations were used (Eq. 4 - 6):

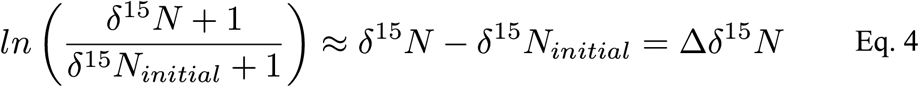

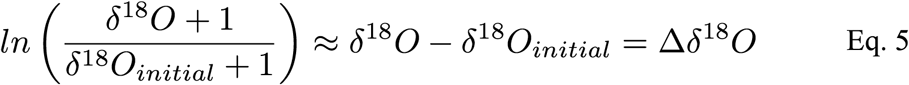

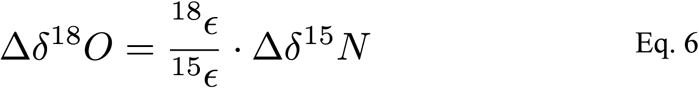

#### Sequence Alignment and Gene Trees

Amino acid sequences for napA and narG reductase genes (see Table S2 for details) were aligned using ClustalOmega Multiple Sequence Alignment ^52^. A list of gene accession numbers is available in SI Table 2. Maximum clade credibility gene trees were constructed using MrBayes’ Markov chain Monte Carlo analysis under an inverse gamma rate variation model with default parameters ^53^.

## Results and Discussion

### Growth of cultures

Growth rates are recorded in SI Table 1. All growth curves and nitrate consumption data are depicted in SI Figs. 1 and 2. No quantitative growth curve data was collected for *S. loihica* and *D. desulfuricans*. While turbidity was detected in *S. loihica*, clumping prevented accurate optical density measurements. *D. desulfuricans* was grown in Postgate’s medium which precipitates iron sulfides and iron hydroxides, preventing accurate optical density measurements. All strains consumed nitrate successfully under fully anaerobic conditions, except for PA14 Δnar which required O_2_ for growth and only consumed significant quantities of nitrate while also exposed to air. Nitrate consumption differed by strain and medium and ranged from as fast as ~15mM nitrate in 8 hours (*B. vireti*) to as slow as 15mM nitrate in 80 hours (PA Δnap). See Fig. S2 for details.

Growth in strains of denitrifying bacteria that cannot perform DNRA (*P. aeruginosa*, *P. denitrificans*) (Fig.1) had little to no nitrite accumulation. However, strains of bacteria that have the potential to perform DNRA in addition to denitrification (*B. vireti*, *B. bataviensis*, *D. desulfuricans*, *S. loihica*) concentrated nitrite during the experiments (SI Fig. 2). This was particularly pronounced in *B. vireti* and *B. bataviensis.* Consequently, later timepoints for these experiments could not be analyzed directly for their nitrate isotopic composition because the sulfamic acid nitrite removal method is only effective to a 7:1 nitrite:nitrate (mol/mol) mixing ratio ^49^. Nitrate in several of these samples with exceedingly high nitrite/ nitrate ratios was thus separated from nitrite by ion chromatography coupled to fraction collection to enable isotopic measurements. The analytical impact of residual nitrite from incomplete nitrite removal by sulfamic acid is discussed in more detail in the SI.

During all time course experiments, decreases in nitrate concentration corresponded to an increasingly enriched residual nitrate pool (SI Figs. 3, 4). Experimental conditions and ^18^ε / ^15^ε proportionality values are summarized in Table 1. ^15^ε values ranged from 10.8 – 34.8‰. ^18^ε values ranged from 5.2 – 29.6‰ (SI Table 1). Isotopic data fit a closed system Rayleigh model for isotope fractionation, with data largely conforming to a linear relationship of δ^15^N or δ^18^O versus the natural logarithm of the remaining nitrate (SI Figs. 3, 4).

**Table 1.** Summary of isotope fractionation experiments. Tracer experiments are included as replicates. Standard error calculated from all experimental replicates.

| Organism | Reductase Gene(s) | Medium | $^{18}\epsilon / ^{15}\epsilon$ +/- std. err. |
| --- | --- | --- | --- |
| *P. aeruginosa* PA14 | both | LB | 0.97 +/- 0.02 |
| *P. aeruginosa* PA14 | both | MOPS | 0.63 +/- 0.02 |
| *P. aeruginosa* $\Delta\text{napA}$ | narG | LB | 0.91 +/- 0.01 |
| *P. aeruginosa* $\Delta\text{napA}$ | narG | MOPS | 0.85 +/- 0.02 |
| *P. aeruginosa* $\Delta\text{narG}$ | napA | LB | 0.49 +/- 0.00 |
| *P. denitrificans* | both | Hahnke | 0.92 +/- 0.01 |
| *B. bataviensis* | narG | TSB | 0.61 +/- 0.06 |
| *B. vireti* | narG | TSB | 0.64 +/- 0.04 |
| *D. desulfuricans* | napA | Postgate | 0.63 +/- 0.06 |
| *S. loihica* | napA | SL | 0.55 +/- 0.01 |

### Nitrate reductases have enzyme specific ^18^ε / ^15^ε coupling

Our data indicate an enzyme specific isotope effect for the Nar and Nap reductases. The PA14 knock-out nitrate reduction experiments show that the Nap reductase in this organism has an ^18^ε / ^15^ε proportionality of 0.49 while that of the Nar reductase in the same organism has a value of 0.86 – 0.91 (Table 1, Fig. 3). The ^18^O tracer experiments confirm that no back reaction of nitrite or exchange with ambient water occurred (SI Fig. 6). The PA14 Δnar data was substantiated by the *D. desulfuricans* and *S. loihica* experiments, with ^18^ε / ^15^ε values of 0.63 +/− 0.06 and 0.55 +/− 0.01, respectively (Table 1; Fig. 4). Together, our data suggest ^18^ε / ^15^ε differences can be purely enzymatic, challenging the hypothesis that environmental ^18^ε / ^15^ε patterns require nitrite re-oxidation from enzymatic reversibility, nitrification or anammox ^27^. These observations for ^18^ε / ^15^ε from nitrate reduction by the Nap reductase in PA14 Δnar are similar to all other available observations from organisms that naturally have only this reductase, with ^18^ε / ^15^ε couplings of 0.63 and 0.51 observed in *R. sphaeroides* and *S. gotlandica*, respectively ^7,31,32^ (Fig. 4).

**Fig. 3.**
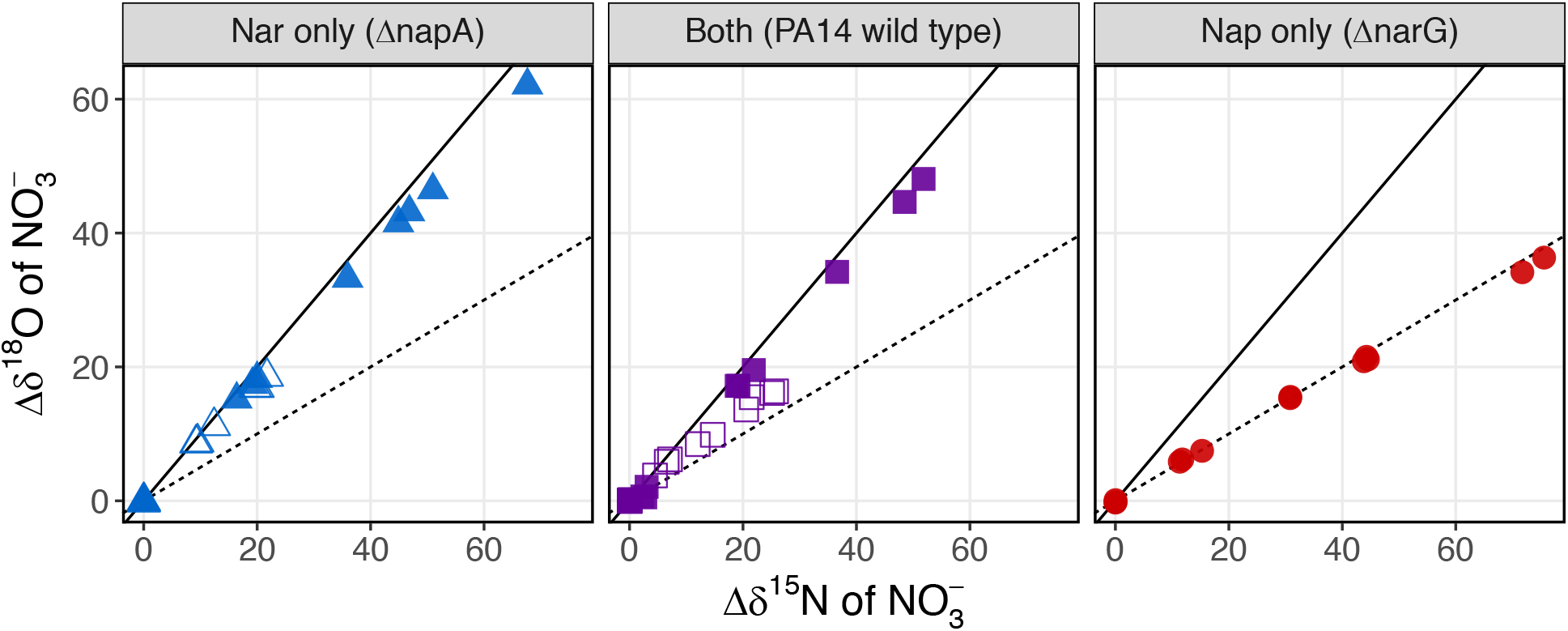
The change in δ^18^O plotted versus change in δ^15^N for the *P. aeruginosa* PA14 wild type (WT) and mutant experiments. “Nar only” corresponds to the PA14 ΔnapA strain, and “Nap only” corresponds to the PA14 ΔnarG strain. Solid lines and dashed lines indicate ^18^ε / ^15^ε proportionalities of 1.0 and 0.5, respectively. Open points indicate cultures grown in MOPS and filled points indicate cultures grown in LB.

**Fig. 4.**
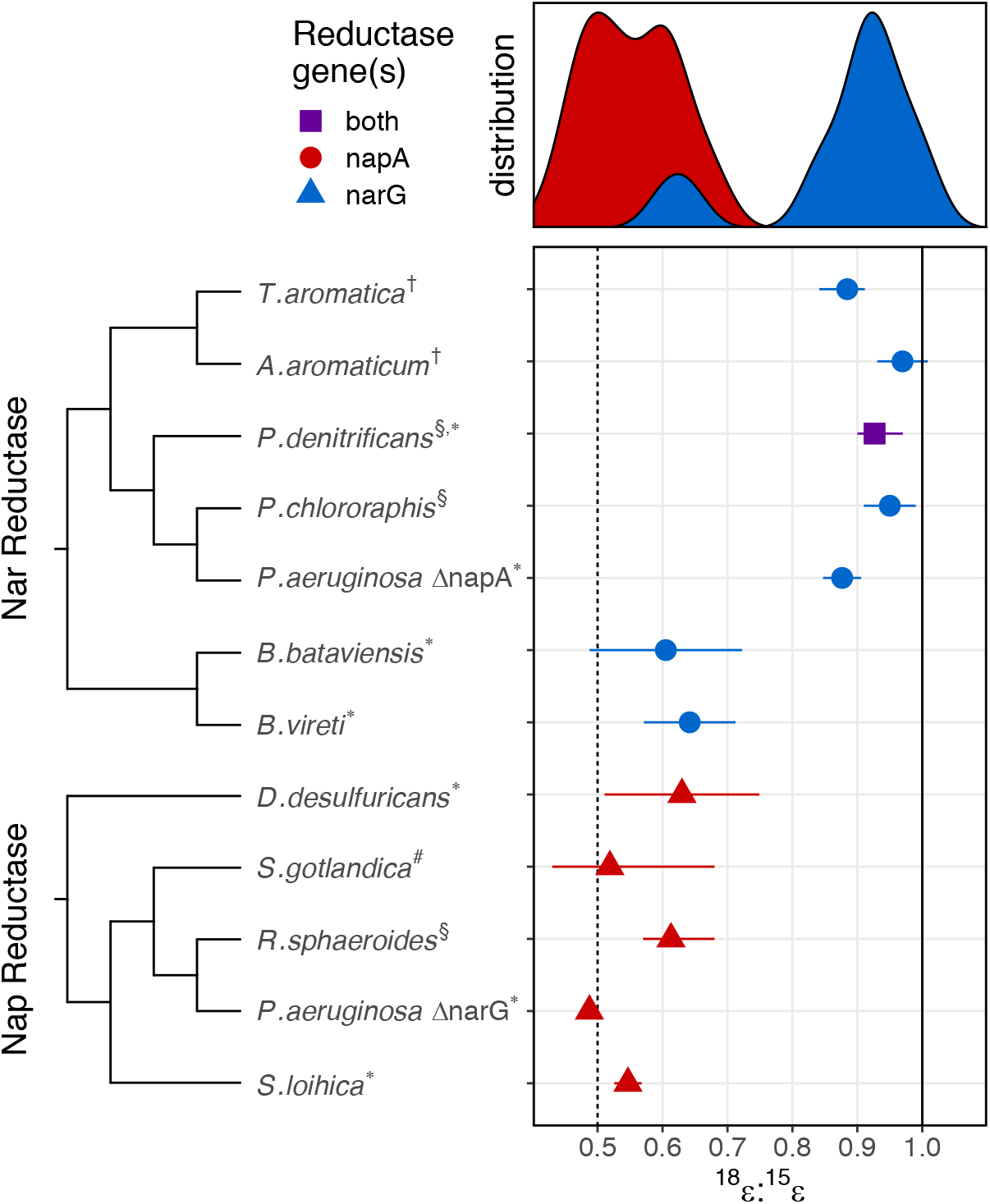
Maximum clade credibility gene trees of the Nap and Nar reductases and a summary of known ^18^ε / ^15^ε values (symbols denote averages, error bars denote value ranges if multiple values available or +/− 2 standard errors for single values) with a distribution of these ranges shown above. Solid line and dashed line indicates ^18^ε / ^15^ε of 1.0 and 0.5, respectively. Colors and shapes indicate the nitrate reductase that is part of the genome of each strain (blue circles: narG only; red triangles: napA only). *P. denitrificans* (purple square) has both genes but under the culturing conditions employed only uses narG ^57,58^. *E. coli* TMAO reductase used as an outgroup in both gene trees. Data collected in this study indicated with an asterisk (*). Literature data collected from (Frey et al., 2014 (#); Granger et al., 2008 (§); Wunderlich et al., 2012 (†)).

As discussed above, the PA14 Δnap strain had an ^18^ε / ^15^ε proportionality of ~0.9 which is consistent with previous reports from organisms that harbor only Nar (Fig. 4) ^7,32–34,54^. Despite having both nitrate reductases present, *P. denitrificans* has been shown in the literature and in our own experiments to also have an ^18^ε / ^15^ε coupling of 0.92 +/− 0.01 ^7,32,33^(Fig. 4). Previous research has demonstrated that *P. denitrificans* PD1222 only uses the Nap reductase under microaerobic conditions and/ or in the presence of highly reduced carbon sources ^55,56^. The culture conditions for *P. denitrificans* used in this study (completely anaerobic conditions, relatively oxidized carbon sources) are not conducive to Nap expression based on the literature data. The ^18^ε / ^15^ε signal we observed in our data is therefore consistent with *P. denitrificans* only reducing nitrate with Nar.

In contrast to all other data on Nar reductases, *B. vireti* and *B. bataviensis* have a significantly lower ^18^ε / ^15^ε: 0.64 +/− 0.04 and 0.61 +/− 0.06, respectively (Table 1; Fig. 4). Although ^18^ε / ^15^ε values between biological replicates covered a wider range than in other organisms, likely due to analytical artifacts from nitrite build-up (see SI discussion on nitrite accumulation), *Bacillus* ^18^ε / ^15^ε values were robustly and consistently lower than all other Nar reductases (Fig. 4). Overall, the *Bacillus* data indicate that it is possible for some Nar reductases to have distinct and lower ^18^ε / ^15^ε proportionality, adding to the complexity of interpreting isotopic signals of nitrate reduction in ecosystems.

### Roles of Nap and Nar reductases

The Nar reductase is known as the primary respiratory reductase amongst denitrifying bacteria. The nar operon is highly conserved, with narGHI present in every known Nar-bearing denitrifier ^28^. Its singular role is in providing energy conservation under anaerobic conditions where high levels of nitrate are present. The Nap reductase, however, has been implicated in both aerobic and traditional anaerobic denitrification, DNRA, redox balancing, nitrate scavenging, and even magnetite biomineralization ^55,56,59–63^. The nap operon is much less conserved, with several combinations of the eleven different genes found across species ^28,30,64,65^. The regulation of these enzymes also differs. As Nar is distinctly used for respiration, the nar operon is upregulated under anaerobic conditions and by the presence of nitrate. Nap regulation, however, is more complicated given the variable operon conformations and assorted functions across species the Nap reductase can perform. For example, reduced carbon sources can upregulate nap expression in some Nap-bearing bacteria ^55,56,58,66^. Additionally, the presence of either oxygen or nitrate can up or down-regulate nap expression depending on species ^63,67–69^. The gene regulation of these enzymes thus ties bacterial preference of reducing nitrate with Nar versus Nap to environmental constraints.

Bacterial preference of using the Nar or Nap reductase was exemplified in the wild type PA14 strain experiments when grown in different mediums. The wild type PA14 strain grown in MOPS medium had an ^18^ε / ^15^ε proportionality of 0.63 +/− 0.02 (Table 1; Fig. 3). This is a midpoint value in comparison to the ^18^ε / ^15^ε proportionality measured in the PA14 Δnap and PA14 Δnar strains and suggests that PA14 was using both nitrate reductases. The Nap reductase for *P. aeruginosa* is used as a backup redox balancing mechanism, in particular under conditions where electron acceptors are limiting ^70^. When grown in LB, this strain exhibited a higher ^18^ε / ^15^ε proportionality of 0.97 +/− 0.02 (Table 1). While LB broth is considered a rich medium, it is actually carbon limited, with mostly amino acids available for uptake ^71^. This would cause lower C/N ratios in contrast to the MOPS minimal medium, in which we provide excess succinate as a carbon source. Past research in *E. coli* has shown that the Nar reductase has a selective advantage under low carbon and high nitrate concentrations, which is the case in our LB grown cultures ^62^. Furthermore, this effect does not occur in the PA14 Δnap strain, suggesting that this is not a difference in how the Nar reductase performs in LB versus minimal medium, but a change in expression pattern by PA14 to maximize energy conservation.

### Mechanism for isotopic differences

Regardless of differences in gene regulation, the Nap and Nar reductases still catalyze the same reaction and yet have different isotope effects. The active site of both reductases are similar, with both containing a Mo-bis-MGD cofactor and iron sulfur cluster. ^28,72^. One distinction is that the Nar reductase’s Mo center is coordinated by an aspartate residue, while the Nap reductase is coordinated by a cysteine. Cysteine is a more reduced residue that may impact the redox potential of the Mo center, affecting how nitrate is bound and reduced ^73–75^. Studies indicate that Nap generally has a higher affinity for nitrate than Nar ^62,76–78^. Furthermore, the base of its substrate channel is lined with positively charged amino acid residues that guides nitrate to the active site ^74,79^. In contrast, Nar has a substrate channel with negatively charged residues that may impact the rate of nitrate binding overall ^80^. Thus, it is possible that the root of isotopic differences lies within the nitrate molecule’s interaction with the active site of these enzymes.

Additionally, it has been proposed that nitrate binds to the catalytic site of Nap and Nar differently. For the Nar reductase, the general mechanism for nitrate binding allows nitrate to bind either Mo(V) or Mo(IV), such that an internal electron transfer may be required before the nitrate molecule can be reduced by Nar ^81,82^. This is in contrast to the Nap reductase where nitrate binds molybdenum only in the reduced state, Mo(IV), and reduces the nitrate immediately ^65,74^. Frey *et al.* (2014) suggested that this may cause a difference in isotope fractionation as the Nar reductase may be subject to an intramolecular isotope effect. While the precise mechanism of nitrate binding and reduction for both Nap and Nar are still uncertain, the Nap reductase’s high affinity for nitrate and its faster reduction mechanism may be key in understanding the differences in ^18^ε / ^15^ε proportionality. Contrary to expectations, our results for the *Bacillus* experiments indicate that a Nap-like isotopic signature with respect to ^18^ε / ^15^ε proportionality is possible in a Nar-reductase. Future work on the structural differences between the *Bacillus* and other Nar reductases may hold the key to uncovering the mechanistic basis for these isotopic differences.

### Interpreting ^18^ε / ^15^ε coupling in ecosystems

Our research shows that nitrate reduction by Nap reductases consistently produces ^18^ε / ^15^ε proportionality values that are lower than those observed in marine ecosystems and may explain the ^18^ε / ^15^ε signals observed in terrestrial ecosystems. The isotopic data sets collected for the terrestrial data in Fig. 2 come from a diverse set of ecosystems ranging from soils to lakes to riparian zones and groundwater runoff from agriculture (see SI for details). Soils in particular can have a large range of redox gradients contained within a few centimeters and experience drastic changes in moisture on short time scales, impacting oxygen availability ^83^. In comparison, marine systems operate at larger scales and experience less heterogeneity over short spatial and temporal scales with dissimilatory nitrate reduction occurring predominately in oxygen minimum zones (OMZ) and anoxic sediments ^14–17,19^. The nar operon has a much narrower regulatory range of permissible environmental conditions than the nap operon and, unlike the latter, is always inhibited by the presence of O_2_ ^84–86^, which may explain the predominance of nar-based nitrate reduction in stable low oxygen systems like OMZs ^87,88^. It is thus conceivable that the Nap reductase’s multiple functions are more suitable for maintaining bacterial homeostasis in terrestrial aquatic ecosystems that can fluctuate significantly over short spatial and temporal timescales.

Though this hypothesis may appear at odds with the established assumption that the Nap reductase is used less commonly than the Nar reductase, limited data is available on Nap versus Nar use in nature. Work by Bru *et al.*^89^ and Smith *et al.*^90^ indicate that Nap and Nar gene copy numbers are roughly equivalent throughout various terrestrial and freshwater environments. Further, slurry incubation experiments performed by Dong *et al.*^91^ indicated that the Nap reductase was more commonly used in one of the three communities of denitrifiers surveyed. While similar studies specifically targeting Nap and Nar gene abundances have not been carried out in marine ecosystems, at minimum this data indicates that the Nap reductase serves an important role in nitrate reduction for bacteria and that its expression is comparable to Nar in freshwater and terrestrial ecosystems.

Since the Nap reductase is not embedded in the cytosolic membrane, and thus not directly involved in proton motive force (PMF) generation, it is frequently presumed to be rarely used for respiration. This explains the common assumption that the isotopic signal of nitrate reduction in ecosystems must stem mainly from the membrane bound cytosolic Nar reductase, as PMF generation is essential for survival and growth ^7,14,17,32,33^. However, the potential to perform nitrate reduction with only a Nap reductase appears to be common place, and with the right auxiliary genes present in the nap operon, can be just as efficient as the Nar reductase at producing a proton motive force (PMF) (Fig. 1B) ^59,65,92^. Future work combining isotopic measurements with quantification of gene expression patterns of the Nap and Nar reductases in different environments can connect our culture-based results back to the trends originally observed in nature. This will be critical when considering the potential impact and extent of *Bacillus*-like Nar enzymes in nature that may have lower ^18^ε / ^15^ε values. The regulation patterns observed in the PA14 wild type strain in MOPS versus LB medium also emphasize the importance of performing transcriptomics over metagenomics, as bacteria with both reductases may switch between Nap and Nar depending on environmental constraints. This is particularly important when considering processes such as DNRA which can use either NapA or NarG to reduce nitrate. Though the Nap reductase is often implicated as the main reductase used during DNRA, many species of bacteria appear to catalyze DNRA solely via the Nar reductase ^43,93–95^. The data presented in this study provides a clear indication that even closely related enzymes can have very distinct isotopic signatures that may allow more comprehensive interpretations of environmental data in the future.

## Acknowledgements

This research was supported by the Department of Geological Sciences at the University of Colorado Boulder and a NASA Exobiology grant (80NSSC17K0667) to SHK. We would like to thank Daniel Sigman for support and access to analytical instrumentation at Princeton University. We are grateful to Emma Kast, Dario Marconi, Sergey Oleynik and other members of the Sigman Lab for providing guidance and support throughout sample processing and isotopic analysis. We also sincerely thank the Dietrich Lab for providing us with the *P. aeruginosa* mutant strains.

