## Supplemental Information for "Enzyme specific isotope effects of the Nap and Nar nitrate reductases"

**Table S1. Data Summary.**

Summary of $^{18}\varepsilon$ / $^{15}\varepsilon$ coupling, fractionation factor, and growth rate estimates by organism and medium. Each row represents a single culture. Fractionation factors calculated from Rayleigh distillation regressions (see Fig. S3 and Fig. S4). Biological replicates with fewer than three isotopic data points were not included. $^{18}\varepsilon$ / $^{15}\varepsilon$ coupling values were calculated using regressions across all cultures from the same medium (see Fig. S5).

| Species | Reductase genes | Medium | $^{18}\varepsilon$ / $^{15}\varepsilon$ | $^{15}\varepsilon$ +/- std. err (‰) | $^{18}\varepsilon$ +/- std. err (‰) | Growth Rate ($hr^{-1}$) +/- std. err |
|---|---|---|---|---|---|---|
| *B. bataviensis* | narG | TSB | 0.61 +/- 0.06 | -15.4 +/- 1.6 | -10.4 +/- 0.7 | 0.31 +/- 0.10 |
| | | | | -13.6 +/- 2.0 | -5.2 +/- 2.1 | 0.24 +/- 0.07 |
| | | | | -11.8 +/- 2.1 | -8.1 +/- 0.7 | 0.24 +/- 0.06 |
| *B. vireti* | narG | TSB | 0.64 +/- 0.04 | -10.8 +/- 3.5 | -9.7 +/- 2.1 | 0.55 +/- 0.05 |
| | | | | -12.6 +/- 5.7 | -9.3 +/- 4.1 | 0.56 +/- 0.04 |
| | | TSB + $^{18}$O water | | -21.3 +/- 3.2 | -11.4 +/- 2.8 | |
| | | | | -21.0 +/- 3.6 | -13.1 +/- 2.2 | |
| | | | | -10.9 +/- 4.6 | -6.3 +/- 2.6 | |
| *D. desulfuricans* | napA | Postgate | 0.63 +/- 0.06 | -22.0 +/- 0.4 | -13.9 +/- 3.1 | |
| | | | | -24.1 +/- 1.2 | -15.2 +/- 2.4 | |
| | | | | -24.8 +/- 1.4 | -15.8 +/- 3.3 | |
| *P. aeruginosa* WT | both | LB | 0.97 +/- 0.02 | -28.4 +/- 1.2 | -26.2 +/- 1.4 | 0.11 +/- 0.04 |
| | | | | -27.7 +/- 0.5 | -25.8 +/- 0.9 | 0.10 +/- 0.04 |
| | | | | -28.5 +/- 0.6 | -26.8 +/- 0.8 | 0.11 +/- 0.03 |
| | | LB + $^{18}$O water | | -22.6 +/- 2.4 | -21.8 +/- 2.2 | 0.05 +/- 0.02 |
| | | | | -26.8 +/- 0.4 | -24.5 +/- 1.3 | 0.06 +/- 0.02 |
| | | | | -23.5 +/- 2.2 | -24.5 +/- 1.4 | 0.09 +/- 0.01 |
| | | MOPS | 0.63 +/- 0.02 | -23.4 +/- 0.9 | -15.3 +/- 1.2 | 0.12 +/- 0.03 |
| | | | | -22.5 +/- 1.0 | -14.3 +/- 1.8 | 0.14 +/- 0.03 |
| | | | | -23.0 +/- 0.4 | -14.1 +/- 1.2 | 0.11 +/- 0.00 |
| *P. aeruginosa* ΔnapA | narG | LB | 0.91 +/- 0.01 | -24.9 +/- 2.0 | -22.7 +/- 1.8 | 0.09 +/- 0.01 |
| | | | | -28.8 +/- 2.9 | -26.2 +/- 2.7 | 0.09 +/- 0.01 |
| | | LB + $^{18}$O water | | -26.7 +/- 0.3 | -24.3 +/- 1.0 | 0.09 +/- 0.01 |
| | | | | -26.5 +/- 0.7 | -24.0 +/- 1.4 | 0.09 +/- 0.01 |
| | | | | -27.5 +/- 0.4 | -23.6 +/- 1.0 | 0.10 +/- 0.01 |
| | | MOPS | 0.85 +/- 0.02 | -26.1 +/- 0.3 | -22.4 +/- 1.2 | 0.09 +/- 0.02 |
| | | | | -24.7 +/- 0.5 | -20.4 +/- 1.6 | 0.07 +/- 0.01 |
| | | | | -23.8 +/- 1.9 | -20.4 +/- 0.7 | 0.08 +/- 0.01 |
| *P. aeruginosa* ΔnarG | napA | LB | 0.49 +/- 0.00 | -31.8 +/- 0.4 | -15.4 +/- 0.4 | 0.46 +/- 0.20 |
| | | | | -32.2 +/- 0.1 | -15.3 +/- 0.0 | 0.41 +/- 0.11 |
| | | | | -33.8 +/- 2.4 | -16.1 +/- 1.6 | 0.39 +/- 0.16 |
| | | LB + $^{18}$O water | | -34.7 +/- 0.8 | -17.4 +/- 0.2 | 0.11 +/- 0.04 |
| | | | | -33.2 +/- 0.5 | -15.7 +/- 0.5 | |
| | | | | -34.8 +/- 0.5 | -17.5 +/- 0.2 | |
| *P. denitrificans* | both | Hahnke | 0.92 +/- 0.01 | -16.9 +/- 1.4 | -15.7 +/- 1.2 | 0.24 +/- 0.04 |
| | | | | -16.7 +/- 0.9 | -15.4 +/- 0.7 | 0.27 +/- 0.01 |
| | | | | -17.0 +/- 0.6 | -15.3 +/- 0.6 | 0.24 +/- 0.02 |
| *S. loihica* | napA | SL | 0.55 +/- 0.01 | -21.8 +/- 0.3 | -12.2 +/- 0.4 | |
| | | | | -20.8 +/- 2.6 | -11.7 +/- 0.2 | |

**Table S1. Gene accession numbers.**

A list of NCBI gene accession numbers used in phylogenetic analyses. In some bacteria two copies of napA exist and thus the analysis was run twice to determine any changes to tree topology. No significant change to the resulting phylogenetic tree occurred. The (*) denotes which gene was used in the final analysis.

| Name | Gene | Gene accession number |
|---|---|---|
| *Aromatoleum aromaticum* EbN1 | Nar | WP_011239378.1 |
| *Paracoccus denitrificans* PD1222 | Nar | WP_011750465.1 |
| *Bacillus bataviensis* LMG 21833 | Nar | EKN65800.1 |
| *Pseudomonas aeruginosa* PA14 | Nar | EOT11604.1 |
| *Bacillus vireti* LMG 21834 | Nar | ETI68959.1 |
| *Pseudomonas chlororaphis chlororaphis* ATCC 9446 | Nar | WP_124303001.1 |
| *Thauera aromatica* K172 | Nar | WP_107220859.1 |
| *Paracoccus denitrificans* PD1222 | Nap | WP_011750941.1 |
| *Rhodobacter sphaeroides* ATCC 17025 | Nap | WP_011910165.1 |
| *Rhodobacter sphaeroides* ATCC 17025 | Nap | WP_011911096.1* |
| *Shewanella loihica* PV-4 | Nap | WP_011867002.1* |
| *Shewanella loihica* PV-4 | Nap | WP_041407111.1 |
| *Sulfurimonas gotlandica* GD1 | Nap | WP_008337904.1 |
| *Desulfovibrio desulfuricans desulfuricans* DSM 642 | Nap | WP_022659785.1 |

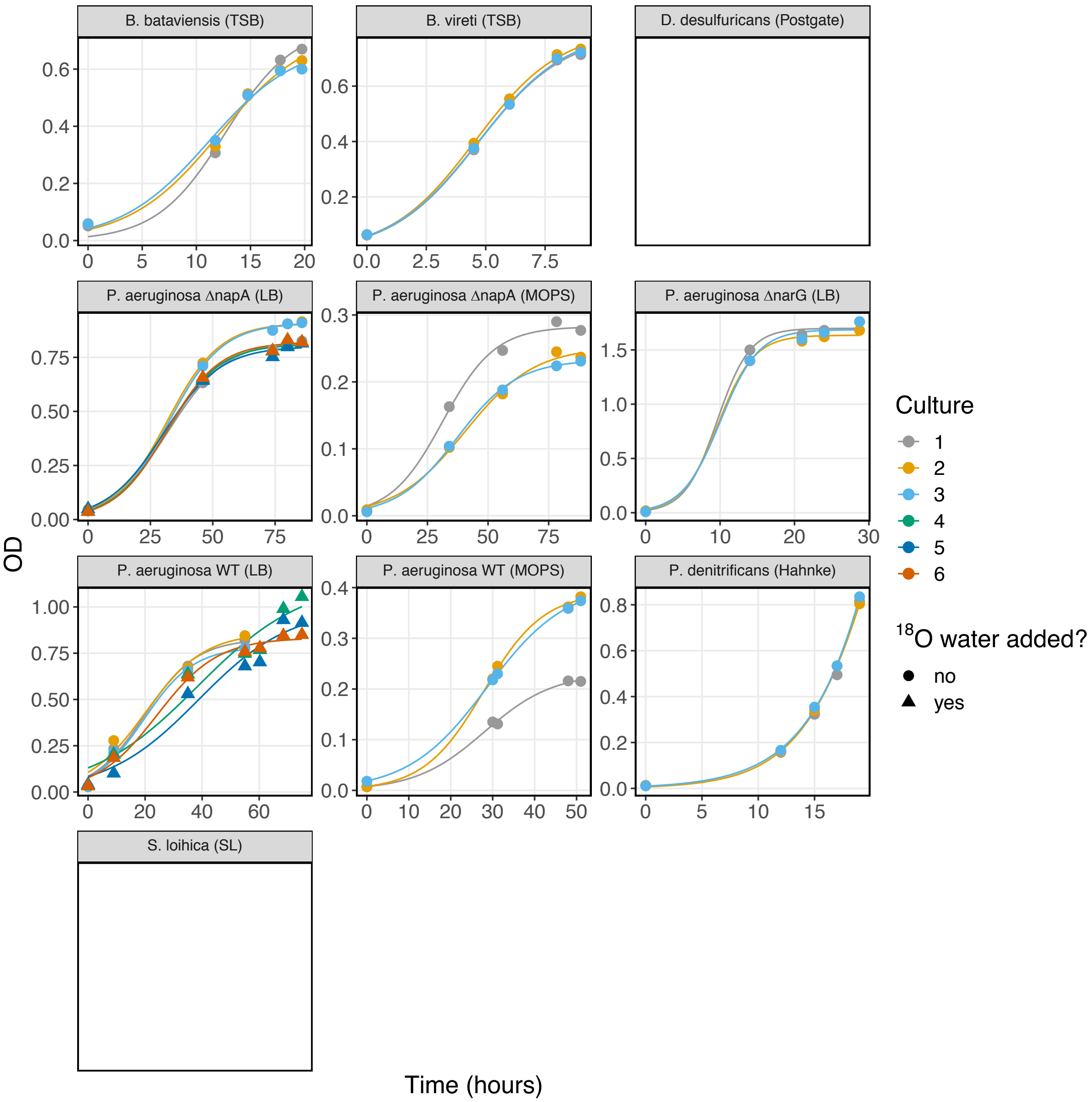


**Figure S1. Growth Curves.**
**Optical density vs. time for all cultures.** Growth curves and parameters were estimated using the R package growthcurver (Sprouffske, 2018) by fitting OD measurements to the following logistic equation, where t is time, OD is the optical density, and fit parameters μ and K represent the growth rate and carrying capacity (max OD), respectively:

$$OD_t = \frac{K}{1 + (K/OD_{t0} - 1) \cdot e^{-\mu t}}$$

No optical density data was collected for *D. desulfuricans* and *S. loihica* which precipitated minerals during growth and could not be measured optically.

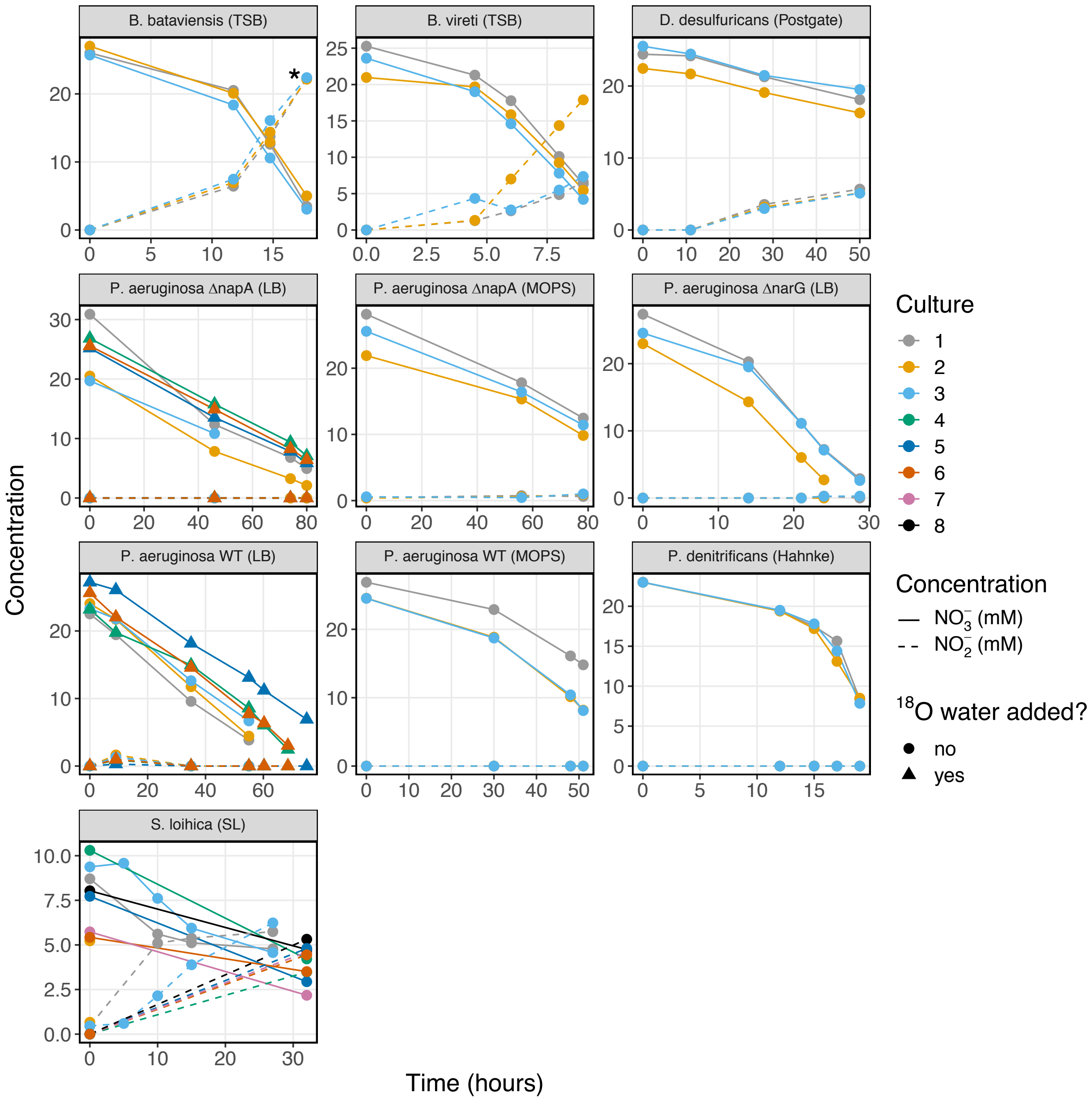


**Figure S2. Nitrate vs time.**
Nitrate and nitrite concentration in each culture versus time. Triangles represent experiments where $^{18}O$ labelled water was added. Starred (*) datapoints indicate samples with > 20mM nitrite accumulation which were purified by ion chromatography prior to analysis (*B. bataviensis* panel).

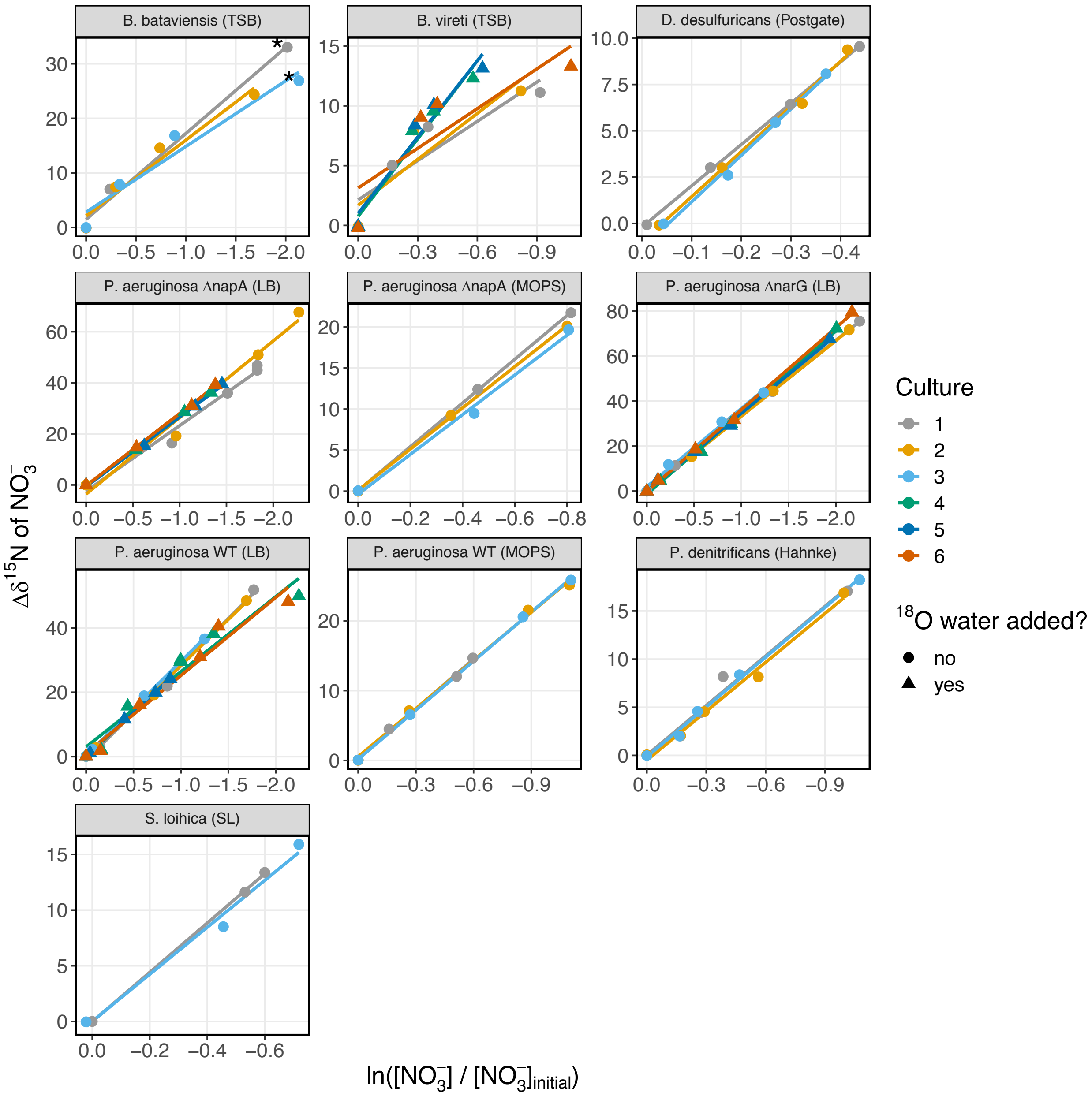


**Figure S3: $\delta^{15}$N vs ln f**

Change in $\delta^{15}$N of nitrate versus the natural log of the remaining nitrate over initial nitrate with linear regression fits to show the isotope fractionation. Triangles represent experiments where $^{18}$O labelled water was added. Starred (*) datapoints indicate samples with > 20mM nitrite accumulation which were purified by ion chromatography prior to analysis (*B. bataviensis* panel).

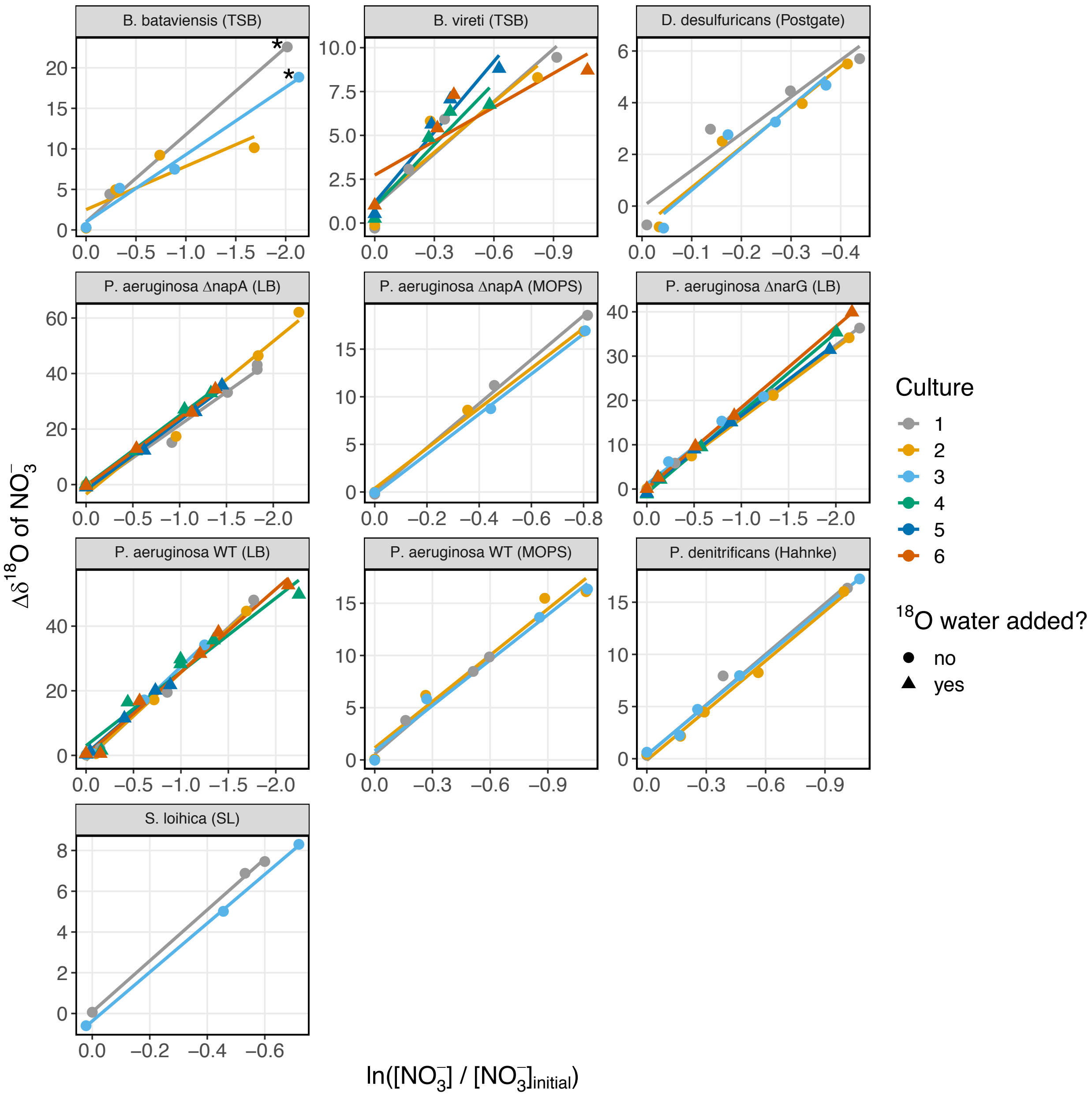


**Figure S4: $\delta^{18}$O vs ln f**

Change in $\delta^{18}$O of nitrate versus the natural log of the remaining nitrate over initial nitrate with linear regression fits to show the isotope fractionation. Triangles represent experiments where $^{18}$O labelled water was added. Starred (*) datapoints indicate samples with > 20mM nitrite accumulation which were purified by ion chromatography prior to analysis (*B. bataviensis* panel).

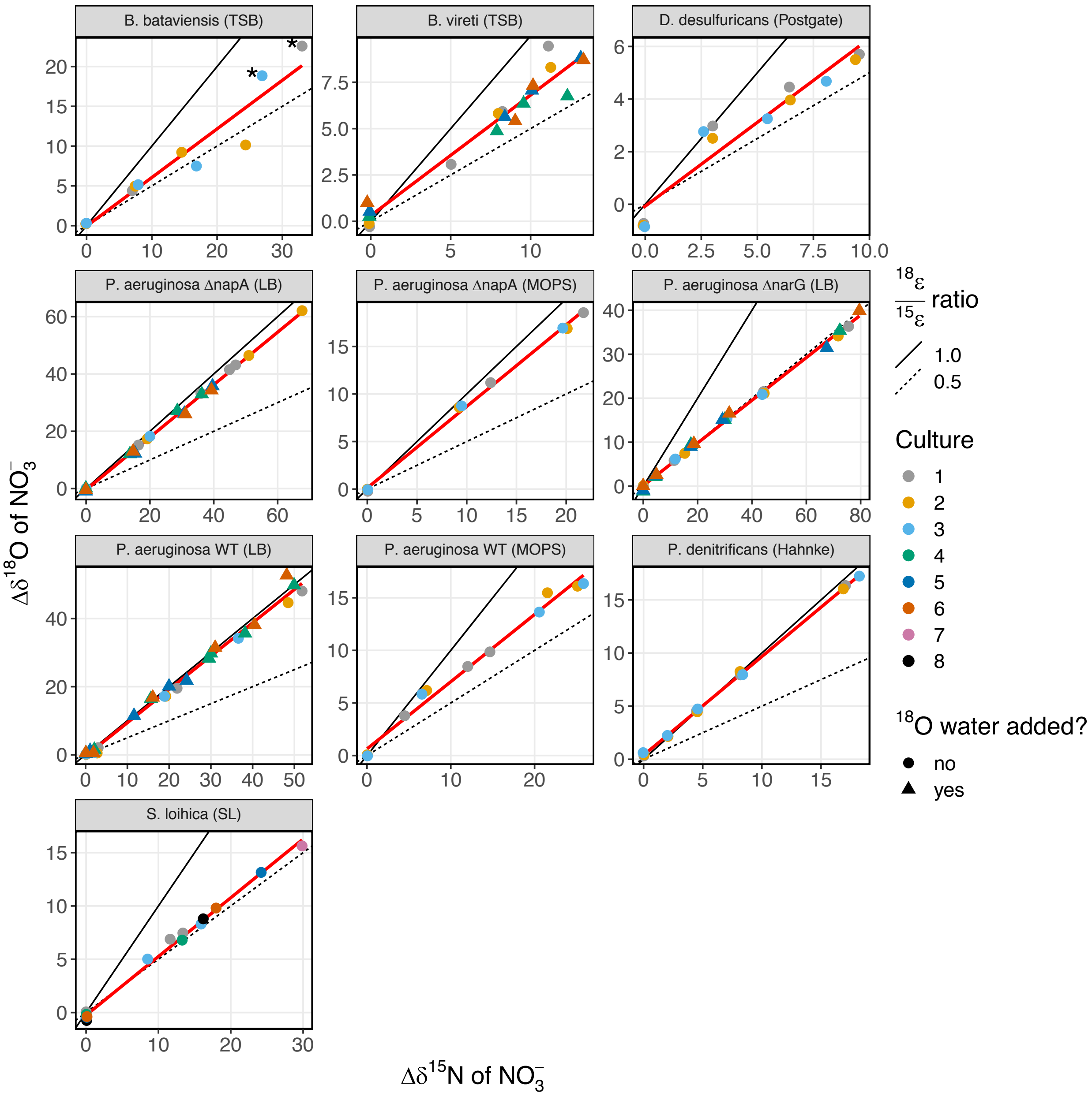


**Figure S5. δ$^{18}$O vs δ$^{15}$N.**
The change in δ$^{18}$O versus change in δ$^{15}$N for all experiments with red regression lines highlighting the $^{18}$ε / $^{15}$ε coupling. Triangles represent experiments where $^{18}$O labelled water was added. Starred (*) datapoints indicate samples with > 20mM nitrite accumulation which were purified by ion chromatography prior to analysis (*B. bataviensis* panel).

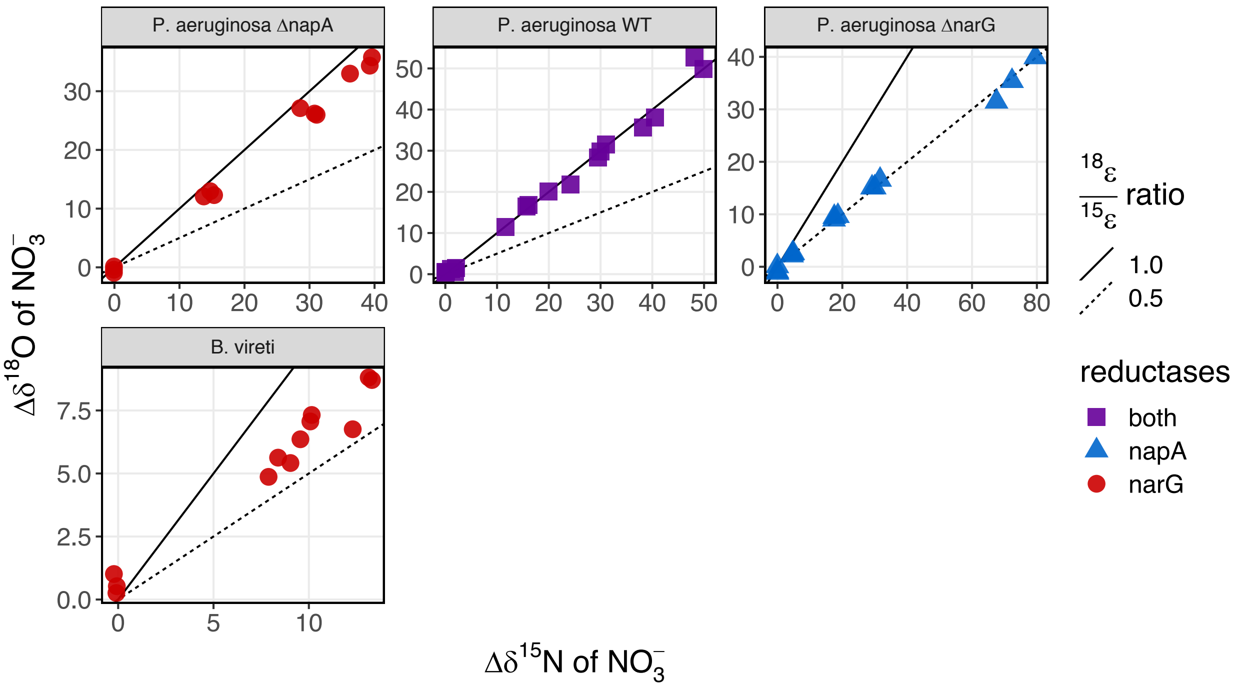


**Figure S6: $^{18}$O water experiments**

The change in $\delta^{18}$O plotted versus the change in $\delta^{15}$N for the PA14 mutant and wild type (WT) and B. vireti tracer experiments. "Nar only" corresponds to the PA Δnap strain, and "Nap only" corresponds to the PA Δnar strain. Solid lines and dashed lines indicate $\varepsilon^{18}$O / $\varepsilon^{15}$N of 1.0 and 0.5, respectively. All PA14 strains grown in LB.

### Supplementary Discussion: the role of nitrite

Nitrite can obscure the experimentally determined oxygen isotope fractionation and $^{18}\varepsilon$ / $^{15}\varepsilon$ coupling associated with nitrate reduction in several ways. First, if nitrite accumulates during a nitrate reduction experiment and is not quantitatively removed prior to nitrate analysis (Granger & Sigman, 2009), it contaminates the nitrate pool and thus affects the resulting isotopic measurement with the denitrifier method (D. M. Sigman et al., 2001). Second, if nitrite is re-oxidized during a nitrate reduction experiment in significant quantities, the kinetic isotope effects of re-oxidation and introduction of water derived oxygen affects the isotopic composition of nitrate (Buchwald & Casciotti, 2010). Both of these scenarios can additionally be affected by abiotic exchange of O between nitrite and water. Below, we discuss how each of these effects would lead to systematic over / under-estimation of the $^{18}\varepsilon$ / $^{15}\varepsilon$ coupling of nitrate reduction and how our experimental results speak to these effects.

#1 Nitrite accumulation and incomplete nitrite removal prior to analysis (= nitrite contamination)

Because nitrate ($NO_3^-$) reduction to nitrite ($NO_2^-$) exhibits normal isotope effects, the residual nitrate becomes enriched in $^{15}$N while the resulting nitrite is depleted in $^{15}$N. However, this simple mass balance consideration does not hold for oxygen because of branching isotope effects. While the residual nitrate does become enriched in $^{18}$O, the resulting nitrite actually has an O isotopic composition similar to the starting nitrate because of the preferential loss of light O to the water (K.L. Casciotti & McIlvin, 2007). In the absence of significant abiotic nitrite-water exchange (see section on exchange below for details), an isotopic measurement of the residual nitrate pool together with some (or all) of this accumulated nitrite using the denitrifier method will lead to a $\delta^{15}N$ measurement that is lower than the actual nitrate pool but a $\delta^{18}O$ measurement that is close to that of the nitrate pool. This would lead to an **overestimate** of the actual $^{18}\varepsilon$ / $^{15}\varepsilon$ coupling (**scenario #1**, **Figure S6**).

#2 Nitrite re-oxidation

During nitrite oxidation, whether mediated by a nitrite oxidoreductase in nitrification or reversibility in the Nar or Nap reductases, two oxygen atoms are inherited from the nitrite molecule being oxidized and the third oxygen atom is derived from water (Buchwald & Casciotti, 2010). In contrast with the N isotopes, the O isotope composition of the resulting nitrate is therefore always affected by the O isotope composition of water ($\delta^{18}O_{H2O}$). For nitrification, there is a normal kinetic isotope effect associated with water incorporation ($^{18}\varepsilon_{k,\,H2O} \approx$ -15‰, (Buchwald & Casciotti, 2010), and an inverse kinetic isotope effect in both O and N associated with the nitrite oxidoreductase ($^{18}\varepsilon_{k,\,NO2}$ = 1.3 – 8.2‰, $^{15}\varepsilon_{k,\,NO2}$ = 12.8‰, (Buchwald & Casciotti, 2010; Karen L. Casciotti, 2009). If nitrite oxidation is quantitative, the kinetic isotope effects of the nitrite oxidoreductase have no effect, but the incorporation of the water O and fractionation associated with it always does. Although the exact isotope effects associated with nitrite oxidation by reversible Nap and Nar reductases are not yet known, it is likely that the contribution of water O in Nap/Nar reversibility would have a similar effect to water O in the nitrite oxidoreductase.

Specifically, because the kinetic isotope affect associated with water incorporation ($^{18}\varepsilon_{k,\,H2O}$) is normal, the O derived from water during nitrite re-oxidation is always isotopically lighter than water (by ≈ - 15‰). If the O isotopic composition of water and starting nitrite/nitrate are comparable, or if water is isotopically lighter than nitrite/nitrate ($\delta^{18}O_{H2O} <= \delta^{18}O_{nitrite} \approx \delta^{18}O_{nitrate}$), nitrite reoxidation would thus contribute isotopically lighter O and lead to an **underestimate** of the $^{18}\varepsilon$ / $^{15}\varepsilon$ coupling of nitrate reduction (**scenario #2, Figure S6**). Only if water is significantly enriched in $^{18}O$ ($\delta^{18}O_{H2O} >> \delta^{18}O_{nitrite} \approx \delta^{18}O_{nitrate}$) could nitrite reoxidation contribute isotopically heavier O and lead to an **overestimate** of the $^{18}\varepsilon$ / $^{15}\varepsilon$ coupling of nitrate reduction (**scenario #3, Figure S6**).

In our experimental conditions with ambient water ($\delta^{18}O_{H2O} \approx$ -16‰, $^{18}O_{nitrate} \approx$ +22‰), nitrite re-oxidation would lead to an underestimate of $^{18}\varepsilon$ / $^{15}\varepsilon$. In our experimental conditions with $^{18}O$ enriched water ($\delta^{18}O_{H2O} \approx$ +100‰, $^{18}O_{nitrate} \approx$ +22‰), nitrite re-oxidation would lead to an overestimate of the $^{18}\varepsilon$ / $^{15}\varepsilon$ coupling.

*O exchange between nitrite and water*

Nitrite can abiotically exchange O with water in a pH-dependent manner on experimentally and environmentally relevant timescales (Buchwald et al., 2012; Buchwald & Casciotti, 2010; Karen L. Casciotti et al., 2010; Grabb et al., 2017). This isotopic equilibration leads to a scrambling of the prior O isotopic signature of nitrite and instead drives it towards water-nitrite equilibrium with fractionation factor $^{18}\varepsilon_{eq} \approx$ 14‰, (K.L. Casciotti & McIlvin, 2007).

In combination with nitrite accumulation and incomplete removal prior to analysis by the denitrifier method (scenario #1 above), water-nitrite equilibration can either counteract the overestimation of the $^{18}\varepsilon$ / $^{15}\varepsilon$ coupling and even lead to an underestimate of $^{18}\varepsilon$ / $^{15}\varepsilon$ if $\delta^{18}O_{H2O}$ + 14‰ < $\delta^{18}O_{nitrate}$ or further exacerbate the overestimation of the $^{18}\varepsilon$ / $^{15}\varepsilon$ if $\delta^{18}O_{H2O}$ + 14‰ > $\delta^{18}O_{nitrate}$. In our experimental conditions with ambient water ($\delta^{18}O_{H2O} \approx$ -16‰, $^{18}O_{nitrate} \approx$ +22‰), this combination would likely lead to no effect or slightly counteracting the underestimate (**scenario #4, Figure S6**). In our experimental conditions with $^{18}O$ enriched water, however ($\delta^{18}O_{H2O} \approx$ +100‰, $^{18}O_{nitrate} \approx$ +22‰), this combination would further **exacerbate the overestimate** of the $^{18}\varepsilon$ / $^{15}\varepsilon$ coupling (**scenario #7, Figure S6**).

On the other hand, water-nitrite equilibration in combination with nitrite re-oxidation (scenario #2 above), can further exacerbate the underestimate of the $^{18}\varepsilon$ / $^{15}\varepsilon$ if $\delta^{18}O_{H2O}$ << $\delta^{18}O_{nitrate}$, counteract the underestimate of the $^{18}\varepsilon$ / $^{15}\varepsilon$ coupling if $\delta^{18}O_{H2O}$ + 14‰ > $\delta^{18}O_{nitrate}$, or lead to an overestimate of the $^{18}\varepsilon$ / $^{15}\varepsilon$ coupling if $\delta^{18}O_{H2O}$ >> $\delta^{18}O_{nitrate}$. In our experimental conditions with ambient water ($\delta^{18}O_{H2O} \approx$ -16‰, $^{18}O_{nitrate} \approx$ +22‰), this combination would further **exacerbate the underestimate** of the $^{18}\varepsilon$ / $^{15}\varepsilon$ coupling (**scenario #5, Figure S6**). In our experimental conditions with $^{18}O$ enriched water ($\delta^{18}O_{H2O} \approx$ +100‰, $^{18}O_{nitrate} \approx$ +22‰), this combination would instead lead to an **overestimate** of the $^{18}\varepsilon$ / $^{15}\varepsilon$ coupling (**scenario #6, Figure S6**).

*Experimental results from experiments with $^{18}O$ enriched water*

If there was significant nitrite re-oxidation, nitrite-reoxidation with water exchange, or nitrite accumulation with nitrite-water exchange occurring in our experimental conditions with $^{18}O$ enriched water, we would expect to observe a significant shift towards higher $^{18}\varepsilon$ / $^{15}\varepsilon$ coupling values compared to the same experiments in ambient water. We did not observe evidence for such a shift in our experiments with $^{18}O$ enriched water, indicating that nitrite re-oxidation and nitrite-water exchange did not play a significant role and did not obscure our experimental determination of the $^{18}\varepsilon$ / $^{15}\varepsilon$ coupling values. However, these experiments cannot rule out that nitrite contamination, *i.e.* nitrite accumulation and incomplete removal prior to analysis (**scenario #1, Fig. S6**) could have occurred under some experimental conditions.

*Nitrite contamination in the B. vireti and B. bataviensis experiments*

The nitrite concentration data from the *Bacillus* strains suggests that significant amounts of nitrite built up over the course of the experiments. While sulfamic acid was used to remove the nitrite, this might have not been quantitative in all samples. Specifically, the non-tracer experiments for *B. vireti* resulted in a higher $^{18}\varepsilon$ / $^{15}\varepsilon$ coupling estimate (0.79 +/- 0.05) than the $^{18}O$ tracer experiments with the same organism (0.58 +/- 0.03). This is the opposite trend we would expect if nitrite re-oxidation or water exchange played a role and suggests that the spread in the data is most likely from variable nitrite accumulation that partly contaminated the nitrate isotope quantification to different degrees. This implies that our estimate for the $^{18}\varepsilon$ / $^{15}\varepsilon$ coupling in *B. veriti* could be an **overestimate** of the true value, i.e. that it could be even more divergent from the expectation for a Nar enzyme (~1.0) than our combined measurements (0.64 +/- 0.04) suggest.

In the second *B. bataviensis* culture we see the final data point decrease in its $^{18}\varepsilon$ / $^{15}\varepsilon$ coupling value. This would suggest that after nitrite accumulation occurred, water then exchanged with the accumulated nitrite. Though sulfamic acid was used to remove nitrite, we see an increase in $\delta^{15}N_{nitrate}$ but almost no change in $\delta^{18}O_{nitrate}$, indicating that nitrite was incompletely removed before sample analysis as discussed in scenario #4. This replicate thus is likely an **underestimate** of the true $^{18}\varepsilon$ / $^{15}\varepsilon$ coupling. However, for *B. bataviensis* cultures one and three, we were able to use ion chromatography to separate nitrite from nitrate with a fraction collector. The final data points for those replicates appear more consistent with the $^{18}\varepsilon$ / $^{15}\varepsilon$ coupling trajectory of earlier datapoints and show no indication of nitrite contamination.

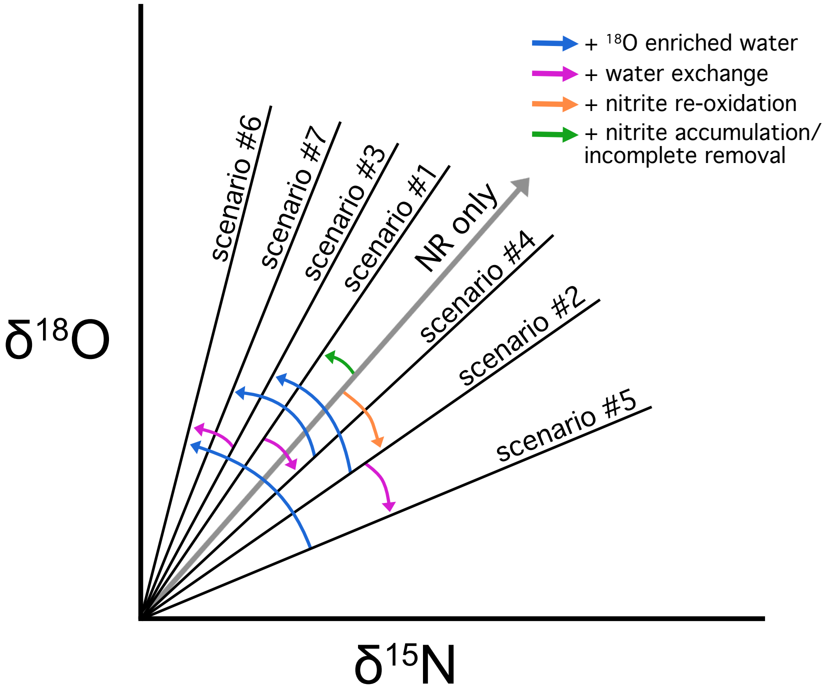


**SI. Fig S6**: A schematic depicting the impact of nitrite isotope effects on $^{18}\varepsilon$ / $^{15}\varepsilon$ values. The grey line represents an $^{18}\varepsilon$ / $^{15}\varepsilon$ value of unidirectional fractionation via nitrate reduction (NR). This represents conditions where only nitrate reduction occurs and there is no exchange with ambient water. Arrows indicate the conditions that change the $^{18}\varepsilon$ / $^{15}\varepsilon$ coupling in each scenario. Scenario #1 represents $^{18}\varepsilon$ / $^{15}\varepsilon$ values where NR occurs with nitrite accumulation and incomplete nitrite removal before analyzing samples. Scenario #2 shows the $^{18}\varepsilon$ / $^{15}\varepsilon$ values where NR occurs in tandem with nitrite re-oxidation with low $\delta^{18}O_{water}$. Scenario #3 shows the impacts of NR with nitrite re-oxidation in elevated $\delta^{18}O_{water}$. Scenario #4 represents NR with nitrite accumulation/ incomplete removal where nitrite-water exchange has occurred. Scenario #5 shows NR with nitrite re-oxidation and nitrite-water exchange in low $\delta^{18}O_{water}$. Scenario #6 shows NR with nitrite re-oxidation and nitrite-water exchange in elevated $\delta^{18}O_{water}$. Scenario #7 shows the impacts of NR with nitrite accumulation and incomplete nitrite removal, in addition to nitrite-water exchange occurring in elevated $\delta^{18}O_{water}$.

*Terrestrial Datasets*

The data displayed in Figure 2 includes terrestrial datasets available in the literature with paired $\delta^{15}N$ and $\delta^{18}O$ measurements from samples collected along transects with likely the same initial nitrate source. Below are descriptions of each dataset and the relevant tables/figures in the original publications.

1. Böttcher *et al.* (1990) measured groundwater samples from different depths in wells impacted by agricultural land use. See $\delta^{15}N$ and $\delta^{18}O$ data in Table 1 and Figure 2 from the arable sampling sites considered by the authors to reflect the same recharge (N5, N10, N11, N12).

2. Aravena & Robertson (1998) measured samples along a groundwater flow path of septic contamination in an aquifer. See $\delta^{15}N$ and $\delta^{18}O$ data in Figure 5.

3. Cey *et al.* (1999) measured groundwater samples along a gradient from agricultural runoff into a riparian zone. See $\delta^{15}N$ and $\delta^{18}O$ data in Figure 12.

4. Mengis *et al.* (1999) measured samples from groundwater along a drainage creek. See $\delta^{15}N$ and $\delta^{18}O$ data in Table 1.

5. Lehmann *et al.* (2003) measured samples from a depth profile through the hypolimnion in the southern basin of Lake Lugano. See $\delta^{15}N$ and $\delta^{18}O$ data in Figure 3.

6. Wenk *et al.* (2014) measured samples from a depth profile through the oxic hypolimnion and the redox transition zone in the northern basin of Lake Lugano. See $\delta^{15}N$ and $\delta^{18}O$ data in Figure 5 (excluding the epilimnion).

Marine Datasets

A much large number of datasets with paired $\delta^{15}N$ and $\delta^{18}O$ measurements exists for marine than for terrestrial samples. The data included in Figure 2 was derived from a compilation by Fripiat et al. (in review) and includes data from the following publications: Bourbonnais et al., 2009; K. L. Casciotti et al., 2018; Karen L. Casciotti et al., 2013; K.L. Casciotti & McIlvin, 2007; Dehairs et al., 2015; DeVries et al., 2013; DiFiore et al., 2009; Gaye et al., 2013; Harms et al., 2019; Kemeny et al., 2016; Knapp et al., 2008, 2011; M. F. Lehmann et al., 2005; N. Lehmann et al., 2018; Marconi et al., 2015; Marconi, Kopf, et al., 2017; Marconi, Sigman, et al., 2017; Martin & Casciotti, 2017; Pantoja et al., 2002; Peng et al., 2018; Rafter et al., 2012, 2013; Rafter & Sigman, 2016; Daniel M. Sigman et al., 2009; Smart et al., 2015; Trull et al., 2008; Van Oostende et al., 2017; Yoshikawa et al., 2018.
